# Effect of gender of new-born, antenatal care and postnatal care on breastfeeding practices in Ethiopia: Evidence from systematic review and meta-analysis of national studies

**DOI:** 10.1101/405605

**Authors:** Tesfa Dejenie Habtewold, Nigussie Tadesse Sharew, Sisay Mulugeta Alemu

## Abstract

**Objective:** The aim of this systematic review and meta-analysis was to investigate the association of gender of new-born, antenatal care (ANC) and postnatal care (PNC) with timely initiation of breastfeeding (TIBF) and exclusive breastfeeding (EBF) practice in Ethiopia.

**Design:** Systematic review and meta-analysis

**Methods:** PubMed, EMBASE, CINAHL, WHO Global Health Library, Web of Science and SCOPUS databases systematically searched and complemented by manual searches to retrieve all available literature. Newcastle-Ottawa Scale (NOS) was used for quality assessment of included studies. Egger’s regression test at p-value threshold ≤ 0.01 was used to examine publication bias. Cochran’s Q X^2^ test, τ^2^, and I^2^ statistics were used to test heterogeneity, estimate amount of total/residual heterogeneity and measure variability attributed to heterogeneity respectively. A meta-analysis using a weighted inverse variance random-effects model was performed. The trend of evidence over time was evaluated by performing a cumulative meta-analysis. Furthermore, mixed-effects meta-regression analysis was done to identify possible sources of heterogeneity.

**Results:** Of 523 articles retrieved, 17 studies (N = 26,146 mothers) on TIBF and 27 studies (N = 17,819 mothers) on EBF were included in the final analysis. ANC (OR = 2.24, 95% CI 1.65 -3.04, p <0.001, I^2^ = 90.9%), PNC (OR = 1.86, 95% CI 1.41 - 2.47, p <0.001, I^2^ = 63.4%) and gender of new-born (OR = 1.31, 95% CI 1.01 - 1.68, p = 0.04, I^2^ = 81.7%) significantly associated with EBF. In addition, ANC (OR = 1.70, 95% CI 1.10 - 2.65, p = 0.02, I^2^ = 93.1%) was significantly associated with TIBF but not gender of new-born (OR = 1.02, 95% CI 0.86 -1.21, p = 0.82, I^2^ = 66.2%).

**Conclusions:** In line with our hypothesis, gender of new-born, ANC and PNC significantly associated with EBF. Likewise, ANC significantly associated with TIBF. Optimal care during pregnancy and after birth is important to ensure adequate breastfeeding. This meta-analysis study provided evidence on breastfeeding practices and its associated factors in Ethiopian context, which can be useful for cross-country and cross-cultural comparison and for breastfeeding improvement initiative in Ethiopia.

**Protocol registration and publication:** <u>CRD42017056768</u> and <u>10.1136/BMJOPEN-2017-017437</u>

**Strengths and limitations of this study:**

- This systematic review and meta-analysis was conducted based on the registered and published protocol.
- Since it is the first study in Ethiopia, the information could be helpful for future researchers, public health practitioners, and healthcare policymakers.
- Almost all included studies were observational which may hinder causality inference.
- Perhaps the results may not be nationally representative given that studies from some regions are lacking.
- Based on the conventional methods of the heterogeneity test, a few analyses suffer from high between-study variation.

## Introduction

For maintaining maternal and new-born health^1^, World Health Organization (WHO) and United Nations Children’s Fund (UNICEF) recommends timely initiation of breastfeeding (TIBF) (i.e. initiating breastfeeding within one hour of birth) and exclusive breastfeeding (EBF) (i.e. feeding only human milk during the first six months).^2^ Breastfeeding provides optimal nutrition for the new-born, increase cognitive development, reduce morbidity and mortality, and preventing newborn and maternal long-term chronic diseases; for example, TIBF prevents 22 % of neonatal deaths.^3^ Inappropriate breastfeeding practice, on the other hand, causes more than two-thirds of under-five child mortality, of which 41% of these deaths occur in Sub-Saharan Africa.^2,4^

According to a new 2017 global report^5^ from the UNICEF and the WHO, only 42%, the majority born in low- and middle-income countries, start breastfeeding within an hour of birth, leaving an estimated 78 million newborns to wait over 1 hour to be put to the breast. The prevalence rate of TIBF varies widely across regions from 35% in the Middle East and North Africa to 65%% in Eastern and Southern Africa. A similar report^6^ also shows that only two in five infants less than 6 months of age are exclusively breastfed. The prevalence rate of EBF ranges from 22% in East Asia and Pacific to 56% in Eastern and Southern Africa.^6^ In 2018, based on our current meta-analysis^7^, the prevalence of TIBF and EBF in Ethiopia is 66.5% and 60.1% respectively. To date, globally, only 22 nations have achieved WHO goal of 70% coverage in TIBF and 23 countries have achieved at least 60% coverage in EBF.^1^

WHO and UNICEF have been working in developing countries for the actualization of optimal breastfeeding and several studies have been conducted on breastfeeding advantages. However, it is challenging to achieve the standard and attributed to several factors including antenatal (ANC), post-natal care (PNC), and gender of new-born^8,9^ and breastfeeding coverage continued to be sub-optimal as a result. In Ethiopia, two meta-analyses studies were conducted.^10,11^ In our previous meta-analysis, we investigated the association between maternal employment, lactation counseling, mode of delivery, place of delivery, maternal age, new-born age and discarding colostrum and, TIBF and EBF.^7,12^ We also investigated whether TIBF associated with EBF.^7^ However, none of these meta-analyses fully studied the effect of gender of new-born, ANC, and PNC. Therefore, we aimed to investigate whether TIBF and EBF in Ethiopia influenced by gender of new-born, ANC and PNC. We hypothesized at least one ANC or PNC visit significantly increase the odds of TIBF and EBF practices. Additionally, mothers with male new-born have higher odds of TIBF and EBF compared to mothers of female newborn.

## Methods

### Protocol registration and publication

The study protocol was registered with the University of York, Centre for Reviews and Dissemination, International prospective register of systematic reviews (PROSPERO) (<u>CRD42017056768</u>) and published.^13^

### Search strategy and databases

PubMed, EMBASE, Cumulative Index to Nursing and Allied Health Literature (CINAHL), WHO Global Health Library, Web of Science and SCOPUS electronic databases were explored to extract all available literature. The search strategy was developed using Population Exposure Controls and Outcome (PECO) searching guide in consultation with a medical information specialist (Supplementary file 1). Searches began 01 August 2017, and the last search was 16 September 2018. Gray literature and cross-references of identified articles and previous meta-analysis were also hand searched.

### PECO guide

> **Population**: All mothers with new-born up to 23 months of age.
>
> **Exposure**: Gender of the new-born, ANC and PNC visit (at least one).
>
> **Controls**: Female new-born, no ANC visit, and no PNC visit.
>
> **Outcome**: TIBF and EBF practice.

### Inclusion and exclusion criteria

Studies were included if they met the following criteria: (1) observational studies including cross-sectional, case-control, cohort studies; (2) conducted in Ethiopia; (3) published in English language; and (4) published between 2000 and 2018. Studies were excluded on any one of the following conditions: (1) study population with HIV/AIDS, preterm new-born and new-born in intensive care unit (ICU); (2) publishing language other than English; (3) abstracts without full-text; and (4) qualitative studies, symposium/conference proceedings, essays, commentaries, editorials and case reports.

### Selection and quality assessment

Initially, all identified articles were exported to Refwork citation manager (RefWorks 2.0; ProQuest LLC, Bethesda, Maryland, USA, http://www.refworks.com) and duplicate studies were canceled. Next, a pair of independent reviewers identified articles by analyzing the abstract and title for relevance and its compliance with the proposed review topic. Agreement between the two reviewers, as measured by Cohen’s Kappa,^14^ was 0.76. After removing irrelevant studies through a respective decision after discussion, full-texts were systematically reviewed for further eligibility analysis. Newcastle-Ottawa Scale (NOS) was used to examine the quality of studies and for potential risk of bias.^15^ In line with the WHO standard definition, outcome measurements were TIBF (the percentage of new-born who breastfeed within the first hour of birth) and EBF (the percentage of infants who exclusively breastfed up to 6 months since birth). Finally, Joanna Briggs Institute (JBI) tool^16^ was used to extract the following data: study area (region and place), method (design), population, number of mothers (calculated sample size and participated in the study) and observed data (i.e. 2 x 2 table). Geographic regions were categorized based on the current Federal Democratic Republic of Ethiopia administrative structure.^17^ Disagreement between reviewers was solved through discussion and consensus.

### Statistical analysis

A meta-analysis using a weighted inverse variance random-effects model was performed to obtain a pooled odds ratio (OR). In addition, to illustrate the trend of evidence regarding the effect of gender of new-born, ANC and PNC on breastfeeding practices, a cumulative meta-analysis was done. Publication bias was assessed by visual inspection of a funnel plot and Egger’s regression test for funnel plot asymmetry using standard error as a predictor in mixed-effects meta-regression model at p-value threshold ≤ 0.01.^18^ Duval and Tweedie trim-and-fill method^19^ was used in case of asymmetric funnel which indicates publication bias. Cochran’s Q X^2^ test, τ^2^, and I^2^ statistics were used to test heterogeneity, estimate amount of total/residual heterogeneity and measure variability attributed to heterogeneity respectively;^20^ for this meta-analysis, we used a reference value of I^2^ > 80% indicating substantial variability related to heterogeneity.^13^ Mixed-effects meta-regression analysis was done to identify possible sources of between-study heterogeneity. The data were analyzed using “metafor” packages in R software version 3.2.1 for Windows.^21^

### Data synthesis and reporting

We analyzed the data in two groups based on TIBF and EBF outcome measurements. Results for each variable were shown using forest plots. Preferred Reporting Items for Systematic Reviews and Meta-Analyses (PRISMA) guidelines for literature review was strictly followed to report our results.^22^

### Minor post hoc protocol changes

Based on the authors’ decision and reviewers recommendation, the following changes were made to our published protocol methods.^13^ We added the Joanna Briggs Institute (JBI) tool^16^ to extract the data. In addition, we used the Duval and Tweedie trim-and-fill method to manage publication bias. Furthermore, cumulative meta-analysis and mixed-effects meta-regression analysis were done to reveal the trend of evidence on each association factor and identify possible sources of between-study heterogeneity respectively.

### Patient and public involvement

The research question and outcome measures were developed by the authors (TD and NT) in consultation with public health professionals and previous studies. Given this is a systematic review and meta-analysis based on published data, patients/study participants were not directly involved in the design and analysis of this study. The results of this study will be disseminated to patients/study participants through health education on factors affecting breastfeeding and disseminating the key findings using brochure in the local language.

## Results

### Search results

In total, we obtained 523 articles from PubMed (n = 169), EMBASE (n = 24), Web of Science (n = 200), SCOPUS (n = 85) and, CINHAL and WHO Global Health Library (n = 5). Fifty additional articles were found through manual search. After removing duplicates and screening of titles and abstracts, 84 studies were selected for full-text review. Forty-three articles were excluded due to several reasons: 19 studies on complementary feeding, 3 studies on pre-lacteal feeding, 3 studies on malnutrition, 17 studies with different variables of interest and one project review report. As a result, 41 articles fulfilled the inclusion criteria and used in this meta-analysis: 17 studies investigated the association between TIBF and gender of new-born and ANC whereas 24 studies between EBF and gender of newborn, ANC, and PNC. The PRISMA flow diagram of literature screening and selection process is shown in figure 1. One study could report more than one outcome measures or associated factors.

**Figure 1:**
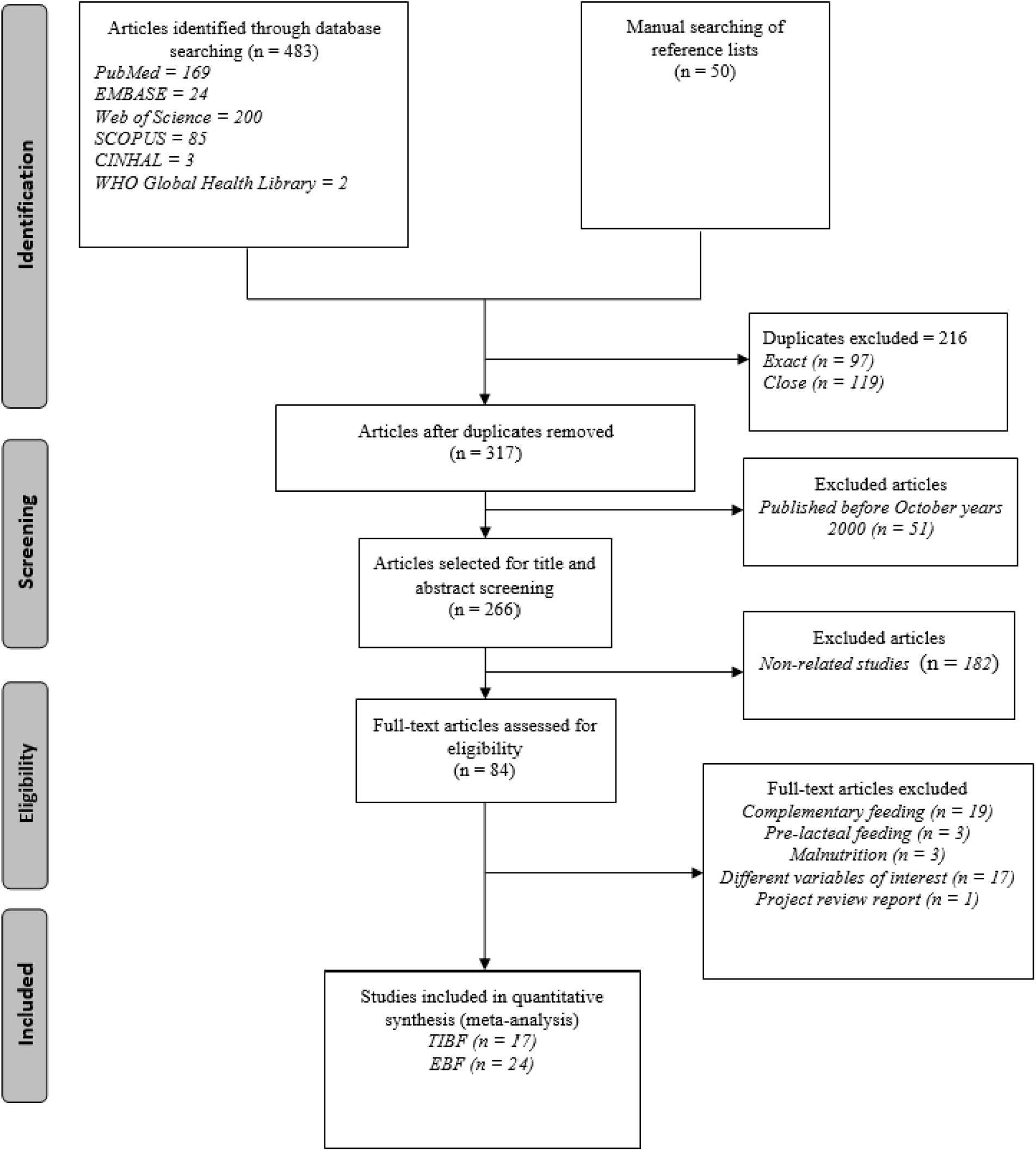
PRISMA flow diagram of literature screening and selection process; “n” in each stage represents the total number of studies that fulfilled particular criteria.

### Study characteristics

As presented in table 1, 17 studies reported the association of TIBF and gender of new-born and ANC in 26,146 mothers. Among these studies, 13 of them were conducted in Amhara (n=5), Oromia (n=4) and Southern Nations, Nationalities and Peoples’ (SNNP) (n=4) region. Regarding residence status of women, eight studies were conducted in both urban and rural whereas six studies in urban dwellers.

**Table 1:** Characteristics of included studies on TIBF.

| Author/publication year | Study area | Study design | Study population | Sample size/ Participated | Factors | TIBF (outcome) |  |  |
| --- | --- | --- | --- | --- | --- | --- | --- | --- |
|  |  |  |  |  |  | Within 1 hour | After 1 hour | Total |
| A. Gender of newborn versus TIBF |  |  |  |  |  |  |  |  |
| Regassa, 2014<sup>23</sup> | SNNPR, Sidama zone | Cross-sectional study | mothers with infants aged between 0 and 6 months old | 1100/ 1094 | Male | 488 | 107 | 595 |
|  |  |  |  |  | Femal | 389 | 110 | 499 |
|  |  |  |  |  | Total | 877 | 217 | 1094 |
| Alemayehu et.al. 2014<sup>24</sup> | Tigray, Axum town | Cross-sectional study | mothers who had children aged 6-12 months | 418/418 | Male | 75 | 141 | 216 |
|  |  |  |  |  | Femal | 99 | 103 | 202 |
|  |  |  |  |  | Total | 174 | 244 | 418 |
| Berhe et.al. 2013<sup>25</sup> | Tigray, Mekelle town | Cross-sectional study | mothers of children aged 0 to 24 months | 361/361 | Male | 166 | 42 | 208 |
|  |  |  |  |  | Femal | 112 | 37 | 149 |
|  |  |  |  |  | Total | 278 | 79 | 357 |
| Beyene et.al. 2017<sup>26</sup> | SNNPR, Dale Woreda | Cross-sectional study | mothers of children under 24 months | 634/ 634 | Male | 262 | 51 | 313 |
|  |  |  |  |  | Femal | 255 | 50 | 305 |
|  |  |  |  |  | Total | 517 | 101 | 618 |
| Lakew et.al. 2015<sup>27</sup> | National | Cross-sectional study * | mothers who had children less than 5 years | 11,654/11,553 | Male | 3124 | 2860 | 5984 |
|  |  |  |  |  | Femal | 3057 | 2511 | 5568 |
|  |  |  |  |  | Total | 6181 | 5371 | 11552 |
| Liben et.al. 2016<sup>28</sup> | Afar, Dubti town | Cross-sectional study | mothers of infants aged less than 6 months | 346/333 | Male | 81 | 122 | 203 |
|  |  |  |  |  | Femal | 70 | 130 | 200 |
|  |  |  |  |  | Total | 151 | 252 | 403 |
| Setegn et.al. 2011<sup>29</sup> | Oromia, Goba district | cross sectional study | mothers with children (< 12 months | 668/ 608 | Male | 164 | 152 | 316 |
|  |  |  |  |  | Femal | 150 | 133 | 283 |
|  |  |  |  |  | Total | 314 | 285 | 599 |
| Wolde et.al. 2014<sup>30</sup> | Oromia, Nekemte town | Cross-sectional study | mothers who had a child less than 24 month | 182/174 | Male | 70 | 10 | 80 |
|  |  |  |  |  | Femal | 84 | 10 | 94 |
|  |  |  |  |  | Total | 154 | 20 | 174 |
| Woldemichael et.al. 2016<sup>31</sup> | Oromia, Tiyo Woreda | Cross-sectional study | mothers who have children less than one year age | 386/373 | Male | 153 | 60 | 213 |
|  |  |  |  |  | Femal | 98 | 62 | 160 |
|  |  |  |  |  | Total | 251 | 122 | 373 |
| Mekonen et.al. | Amhara, | Cross- | mothers of infants | 845/823 | Male | 214 | 229 | 443 |
| 2018<sup>32</sup> | South Gondar | sectional study | under 12 months |  | Female | 187 | 193 | 380 |
|  |  |  |  |  | Total | 401 | 422 | 823 |
| B. Antenatal care versus TIBF |  |  |  |  |  |  |  |  |
| Gultie et.al 2016<sup>33</sup> | Amhara, Debre Berhan town | Cross-sectional study | mothers having children aged less than 23 months old | 548/548 | ANC | 482 | 88 | 570 |
|  |  |  |  |  | No ANC | 16 | 15 | 31 |
|  |  |  |  |  | Total | 498 | 103 | 601 |
| Tamiru et.al 2012<sup>34</sup> | Oromia, Jimma Arjo Woreda | Cross-sectional study | mothers of index children aged 0 to 6 months | 384/ 382 | ANC | 115 | 69 | 184 |
|  |  |  |  |  | No ANC | 120 | 71 | 191 |
|  |  |  |  |  | Total | 235 | 140 | 375 |
| Tamiru et.al 2015<sup>35</sup> | SNNPR, Arba Minch Zuria Woreda | cross-sectional study | mothers of infants aged two years and younger | 384/384 | ANC | 179 | 140 | 319 |
|  |  |  |  |  | No ANC | 40 | 24 | 64 |
|  |  |  |  |  | Total | 219 | 164 | 383 |
| Berhe et.al 2013<sup>25</sup> | Tigray, Mekelle town | Cross-sectional study | mothers of children aged 0 to 24 months | 361/361 | ANC | 263 | 66 | 329 |
|  |  |  |  |  | No ANC | 15 | 13 | 28 |
|  |  |  |  |  | Total | 278 | 79 | 357 |
| Adugna et.al 2014<sup>36</sup> | SNNPR, Arba Minch Zuria | cross-sectional study | women who had children under two years | 384/383 | ANC | 179 | 140 | 319 |
|  |  |  |  |  | No ANC | 40 | 24 | 64 |
|  |  |  |  |  | Total | 219 | 164 | 383 |
| Beyene et.al 2017<sup>26</sup> | SNNPR, Dale Woreda | Cross-sectional study | mothers of children under 24 months | 634/ 634 | ANC | 206 | 58 | 264 |
|  |  |  |  |  | No ANC | 311 | 43 | 354 |
|  |  |  |  |  | Total | 517 | 101 | 618 |
| Derso et.al 2017<sup>37</sup> | Amhara, Dabat district | Cross-sectional study * | mothers with children under five years of age | 6,761/ 6,761 | ANC | 2135 | 2220 | 4355 |
|  |  |  |  |  | No ANC | 670 | 1364 | 2034 |
|  |  |  |  |  | Total | 2805 | 3584 | 6389 |
| Liben et.al 2016<sup>28</sup> | Afar, Dubti town | Cross-sectional study | mothers of infants aged less than 6 months | 346/333 | ANC | 110 | 196 | 306 |
|  |  |  |  |  | No ANC | 41 | 56 | 97 |
|  |  |  |  |  | Total | 151 | 252 | 403 |
| Seid et.al 2013<sup>38</sup> | Amhara, Bahir Dar city | Cross-sectional study | mothers who Delivered in the last 12 months | 819/819 | ANC | 680 | 94 | 774 |
|  |  |  |  |  | No ANC | 29 | 12 | 41 |
|  |  |  |  |  | Total | 709 | 106 | 815 |
| Setegn et.al 2011<sup>29</sup> | Oromia, Goba district | cross sectional study | mothers with children (< 12 months | 668/ 608 | ANC | 270 | 238 | 508 |
|  |  |  |  |  | No ANC | 37 | 19 | 56 |
|  |  |  |  |  | Total | 307 | 257 | 564 |
| Tewabe 2016<sup>39</sup> | Amhara, Motta town | cross-sectional study | mothers with infant less than six month old | 423/405 | ANC | 282 | 41 | 323 |
|  |  |  |  |  | No ANC | 37 | 45 | 82 |
|  |  |  |  |  | Total | 319 | 86 | 405 |
| Woldemichael et.al 2016<sup>31</sup> | Oromia, Tiyo Woreda | Cross-sectional study | mothers who have children Less Than One Year Age | 386/373 | ANC | 194 | 41 | 235 |
|  |  |  |  |  | No ANC | 57 | 81 | 138 |
|  |  |  |  |  | Total | 251 | 122 | 373 |
| Mekonen et.al. 2018<sup>32</sup> | Amhara, South Gondar | Cross-sectional study | mothers of infants under 12 months | 845/823 | ANC | 370 | 332 | 702 |
|  |  |  |  |  | No ANC | 31 | 90 | 121 |
|  |  |  |  |  | Total | 401 | 422 | 823 |

\*= Used nationally representative Ethiopian Demographic Health Survey (EDHS) data

Twenty-four studies reported the association of EBF with gender of new-born, ANC and PNC in 17,819 mothers. Of these studies, 11 were conducted in Amhara and seven in SNNP region. Based on the residence status of women, 10 studies were conducted in urban, eight in urban and rural, and six in rural dwellers. Even though almost all studies were cross-sectional, five studies have used a nationally representative data of the Ethiopian Demographic Health Survey (EDHS) [19-23]. Detailed study characteristics have shown in Table 2.

**Table 2:** Characteristics of included studies on EBF.

| Author/publication year | Study area | Study design | Study population | Sample size/Participated | Factors | EBF (outcome) |  |  |
| --- | --- | --- | --- | --- | --- | --- | --- | --- |
|  |  |  |  |  |  | Yes | No | Total |
| A. Gender of newborn versus EBF |  |  |  |  |  |  |  |  |
| Asemahagn 2016<sup>40</sup> | Amhara, Azezo district | Cross-sectional study | Women having children aged from 0–6 months | 346/ 332 | Male | 95 | 38 | 133 |
|  |  |  |  |  | Femal | 167 | 32 | 199 |
|  |  |  |  |  | Total | 262 | 70 | 332 |
| Setegn et.al. 2012<sup>41</sup> | Oromia, Bale Zone, Goba district | Cross-sectional study | Mothers-infant pairs | 668/608 | Male | 107 | 43 | 150 |
|  |  |  |  |  | Femal | 92 | 37 | 129 |
|  |  |  |  |  | Total | 199 | 80 | 279 |
| Sonko et.al. 2015<sup>42</sup> | SNNPR, Halaba special woreda | Cross-sectional study | Mothers With children less than six months of age | 422/420 | Male | 145 | 60 | 205 |
|  |  |  |  |  | Femal | 151 | 64 | 215 |
|  |  |  |  |  | Total | 296 | 124 | 420 |
| Regassa 2014<sup>23</sup> | SNNPR, Sidama zone | Cross-sectional study | with infants aged between 0 and 6 months old | 1100/ 1094 | Male | 109 | 19 | 128 |
|  |  |  |  |  | Femal | 89 | 17 | 106 |
|  |  |  |  |  | Total | 198 | 36 | 234 |
| Alemayehu et.al. 2014<sup>24</sup> | Tigray, Axum town | Cross-sectional study | mothers who had children aged 6-12 months | 418/418 | Male | 97 | 119 | 216 |
|  |  |  |  |  | Femal | 77 | 128 | 205 |
|  |  |  |  |  | Total | 174 | 247 | 421 |
| Biks et.al. 2015<sup>43</sup> | Amhara, Dabat district | Nested case–control study * | All pregnant women in the second/third trimester | 1,769/1,769 | Male | 271 | 619 | 890 |
|  |  |  |  |  | Femal | 727 | 1148 | 1875 |
|  |  |  |  |  | Total | 998 | 1767 | 2765 |
| Arage et.al. 2016<sup>44</sup> | Amhara, Debre Tabor Town | Cross-sectional study | Mothers of Infants Less Than Six Months of Age | 470/453 | Male | 119 | 40 | 159 |
|  |  |  |  |  | Femal | 227 | 67 | 294 |
|  |  |  |  |  | Total | 346 | 107 | 453 |
| Adugna et.al. 2017<sup>45</sup> | SNNPR, Hawassa city | Cross-sectional study | Mothers with infants aged 0–6 months | 541/529 | Male | 169 | 88 | 257 |
|  |  |  |  |  | Femal | 153 | 119 | 272 |
|  |  |  |  |  | Total | 322 | 207 | 529 |
| Egata et.al. 2013<sup>46</sup> | Oromia, Kersa district | Cross-sectional study* | Mothers of children under-two years of age | 881/860 | Male | 323 | 124 | 447 |
|  |  |  |  |  | Femal | 294 | 119 | 413 |
|  |  |  |  |  | Total | 617 | 243 | 860 |
| Teka et al. 2015<sup>47</sup> | Tigray, | Cross- | Mothers having | 541/530 | Male | 158 | 60 | 218 |
|  | Enderta Woreda | sectional study | children aged less than 24 months |  | Femal | 214 | 98 | 312 |
|  |  |  |  |  | Total | 372 | 158 | 530 |
| Sefene et al. 2013<sup>48</sup> | Amhara, Bahir Dar city | Cross-sectional study | Mothers who had a child age less than 6 months | 170/159 | Male | 36 | 47 | 83 |
|  |  |  |  |  | Femal | 42 | 34 | 76 |
|  |  |  |  |  | Total | 78 | 81 | 159 |
| **B. Antenatal care versus EBF** |  |  |  |  |  |  |  |  |
| Asemahagn 2016<sup>40</sup> | Amhara, Azezo district | Cross-sectional study | Women having children aged from 0–6 months | 346/ 332 | ANC | 243 | 57 | 300 |
|  |  |  |  |  | No ANC | 19 | 13 | 32 |
|  |  |  |  |  | Total | 262 | 70 | 332 |
| Gultie et.al 2016<sup>33</sup> | Amhara, Debre Berhan town | Cross-sectional study | mothers having children aged less than 23 months old | 548/548 | ANC | 263 | 253 | 516 |
|  |  |  |  |  | No ANC | 10 | 21 | 31 |
|  |  |  |  |  | Total | 273 | 274 | 547 |
| Hunegnaw et.al. 2017<sup>49</sup> | Amhara, Gozamin district | Cross-sectional study | Mothers who had Infants aged between 6 and 12 months | 506/478 | ANC | 341 | 109 | 450 |
|  |  |  |  |  | No ANC | 17 | 11 | 28 |
|  |  |  |  |  | Total | 358 | 120 | 478 |
| Lenja et.al. 2016<sup>50</sup> | SNNPR, Offa district | Cross-sectional study | Mothers of infants younger than 6 months | 403/396 | ANC | 233 | 43 | 276 |
|  |  |  |  |  | No ANC | 44 | 88 | 132 |
|  |  |  |  |  | Total | 277 | 131 | 408 |
| Seid et.al 2013<sup>38</sup> | Amhara, Bahir Dar city | Cross-sectional study | Mothers who Delivered in the last 12 months | 819/819 | ANC | 405 | 372 | 777 |
|  |  |  |  |  | No ANC | 7 | 35 | 42 |
|  |  |  |  |  | Total | 412 | 407 | 819 |
| Setegn et.al 2011<sup>29</sup> | Oromia, Goba district | cross sectional study | mothers with children (< 12 months | 668/ 608 | ANC | 166 | 65 | 231 |
|  |  |  |  |  | No ANC | 27 | 10 | 37 |
|  |  |  |  |  | Total | 193 | 75 | 268 |
| Sonko et.al. 2015<sup>42</sup> | SNNPR, Halaba special woreda | Cross-sectional study | Mothers With children less than six months of age | 422/420 | ANC | 258 | 88 | 346 |
|  |  |  |  |  | No ANC | 38 | 36 | 74 |
|  |  |  |  |  | Total | 296 | 124 | 420 |
| Tadesse et.al. 2016<sup>51</sup> | SNNPR, Sorro District | Cross-sectional Study | Mothers With infants aged of 0–5 months | 602/ 579 | ANC | 211 | 121 | 332 |
|  |  |  |  |  | No ANC | 59 | 123 | 182 |
|  |  |  |  |  | Total | 270 | 244 | 514 |
| Tariku et.al. 2017<sup>52</sup> | Amhara, Dabat District | Cross-sectional study * | Mothers with children aged less than 59 months | 5,227/ 5,227 | ANC | 1979 | 1353 | 3332 |
|  |  |  |  |  | No ANC | 713 | 876 | 1589 |
|  |  |  |  |  | Total | 2692 | 2229 | 4921 |
| Tewabe et.al. 2017<sup>39</sup> | Amhara, Motta town, East Gojjam zone | Cross-sectional Study | Mothers with infant less than six months old | 423/405 | ANC | 185 | 164 | 349 |
|  |  |  |  |  | No ANC | 18 | 38 | 56 |
|  |  |  |  |  | Total | 203 | 202 | 405 |
| Tamiru et.al 2012<sup>34</sup> | Oromia, Jimma Arjo Woreda | Cross-sectional study | Mothers of index children aged 0 to 6 months | 384/ 382 | ANC | 87 | 103 | 190 |
|  |  |  |  |  | No ANC | 96 | 96 | 192 |
|  |  |  |  |  | Total | 183 | 199 | 382 |
| Tamiru et.al 2015<sup>35</sup> | SNNPR, Arba Minch Zuria Woreda | cross-sectional study | Mothers of infants aged two years and younger | 384/384 | ANC | 228 | 92 | 320 |
|  |  |  |  |  | No ANC | 27 | 37 | 64 |
|  |  |  |  |  | Total | 255 | 129 | 384 |
| Biks et.al. 2015<sup>43</sup> | Amhara, Dabat | Nested case–control study | All pregnant women in the | 1,769/1,769 | ANC | 180 | 277 | 457 |
|  |  |  |  |  | No ANC | 363 | 949 | 1312 |
|  | district | * | second/third trimester |  | Total | 543 | 1226 | 1769 |
| Abera 2012<sup>53</sup> | Harari, Harar town | Cross-sectional study | Mothers of children aged less than two years | 604/583 | ANC | 194 | 163 | 357 |
|  |  |  |  |  | No ANC | 13 | 29 | 42 |
|  |  |  |  |  | Total | 207 | 192 | 399 |
| Arage et.al. 2016<sup>44</sup> | Amhara, Debre Tabor Town | Cross-sectional study | Mothers of Infants Less Than Six Months of Age | 470/453 | ANC | 384 | 39 | 423 |
|  |  |  |  |  | No ANC | 18 | 12 | 30 |
|  |  |  |  |  | Total | 402 | 51 | 453 |
| Adugna et.al. 2017<sup>45</sup> | SNNPR, Hawassa city | Cross-sectional study | Mothers with infants aged 0–6 months | 541/529 | ANC | 221 | 111 | 332 |
|  |  |  |  |  | No ANC | 101 | 96 | 197 |
|  |  |  |  |  | Total | 322 | 207 | 529 |
| Egata et.al. 2013<sup>46</sup> | Oromia, Kersa district | Cross-sectional study * | Mothers of children under two years of age | 881/860 | ANC | 233 | 135 | 368 |
|  |  |  |  |  | No ANC | 384 | 108 | 492 |
|  |  |  |  |  | Total | 617 | 243 | 860 |
| Taddele et.al. 2014<sup>54</sup> | Amhara, Injibara Town | Comparative cross-sectional study | Employed and unemployed mothers of children age ≤ 1 year | 524/473 | ANC | 90 | 98 | 188 |
|  |  |  |  |  | No ANC | 6 | 23 | 29 |
|  |  |  |  |  | Total | 96 | 121 | 217 |
| Echamo. 2012<sup>55</sup> | SNNPR, Arbaminch town | Cross-sectional study | Mothers of infants within the age of six to twelve months | 768/768 | ANC | 332 | 360 | 692 |
|  |  |  |  |  | No ANC | 25 | 51 | 76 |
|  |  |  |  |  | Total | 357 | 411 | 768 |
| Teka et al. 2015<sup>47</sup> | Tigray, Enderta Woreda | Cross-sectional study | Mothers having children aged less than 24 months | 541/530 | ANC | 325 | 134 | 459 |
|  |  |  |  |  | No ANC | 47 | 24 | 71 |
|  |  |  |  |  | Total | 372 | 158 | 530 |
| Chekol et al. 2017<sup>56</sup> | Amhara, Gondar town | Cross-sectional study | Mothers with children age 7–12 months | 333/333 | ANC | 131 | 117 | 248 |
|  |  |  |  |  | No ANC | 29 | 56 | 85 |
|  |  |  |  |  | Total | 160 | 173 | 333 |
| C. Postnatal care versus EBF |  |  |  |  |  |  |  |  |
| Asemahagn 2016<sup>40</sup> | Amhara, Azezo district | Cross-sectional study | Women having children aged from 0–6 months | 346/ 332 | PNC | 137 | 25 | 162 |
|  |  |  |  |  | No PNC | 125 | 45 | 170 |
|  |  |  |  |  | Total | 262 | 70 | 332 |
| Lenja et.al. 2016<sup>50</sup> | SNNPR, Offa district | Cross-sectional study | Mothers of infants younger than 6 months | 403/396 | PNC | 188 | 33 | 221 |
|  |  |  |  |  | No PNC | 121 | 54 | 175 |
|  |  |  |  |  | Total | 309 | 87 | 396 |
| Sonko et.al. 2015<sup>42</sup> | SNNPR, Halaba special woreda | Cross-sectional study | Mothers with children less than six months of age | 422/420 | PNC | 98 | 25 | 123 |
|  |  |  |  |  | No PNC | 197 | 99 | 296 |
|  |  |  |  |  | Total | 295 | 124 | 419 |
| Tadesse et.al. 2016<sup>51</sup> | SNNPR, Sorro District | Cross-sectional Study | Mothers with infants aged 0–5 months | 602/ 579 | PNC | 204 | 127 | 331 |
|  |  |  |  |  | No PNC | 66 | 117 | 183 |
|  |  |  |  |  | Total | 270 | 244 | 514 |
| Tewabe et.al. 2017<sup>57</sup> | Amhara, Motta town, East Gojjam zone | Cross-sectional Study | Mothers with infant less than six Months old | 423/405 | PNC | 116 | 81 | 197 |
|  |  |  |  |  | No PNC | 87 | 121 | 208 |
|  |  |  |  |  | Total | 203 | 202 | 405 |
| Abera 2012<sup>53</sup> | Harari, Harar town | Cross-sectional | Mothers of children aged less | 604/583 | PNC | 29 | 31 | 60 |
|  |  |  |  |  | No PNC | 178 | 161 | 339 |
|  |  | study | than two years |  | Total | 207 | 192 | 399 |
| Teka et al. 2015<sup>47</sup> | Tigray, Enderta woreda | Cross-sectional study | Mothers having children aged less than 24 months | 541/530 | PNC | 167 | 86 | 253 |
|  |  |  |  |  | No PNC | 205 | 72 | 277 |
|  |  |  |  |  | Total | 372 | 158 | 530 |

\*= Used nationally representative Ethiopian Demographic Health Survey (EDHS) data

### Meta-analysis

#### TIBF

Among the 17 selected studies, 10 studies^23–32^ reported the association between TIBF and gender of new-born in 16,411 mothers (Table 1A). The pooled odds ratio (OR) of gender of new-born was 1.02 (95% CI 0.86 - 1.21, p = 0.82, I^2^ = 66.2%) (figure 2). Mothers with male new-born had 2% higher chance of initiating breastfeeding within one hour of birth compared with female newborn although not statistically significant. Egger’s regression test for funnel plot asymmetry was not significant (z = 0.41, p = 0.68) (Supplementary figure 1).

**Figure 2:**
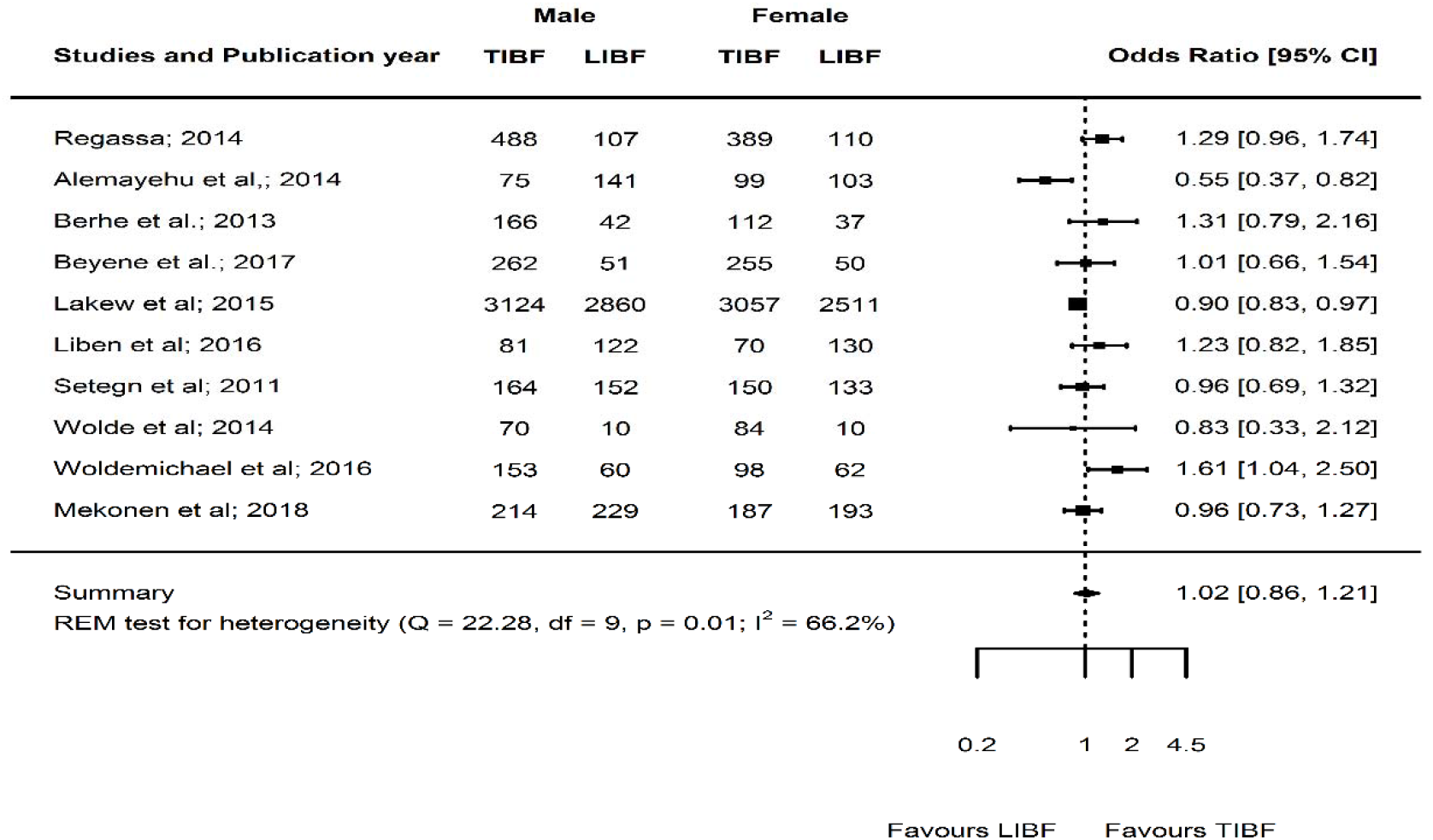
Forest plot of the unadjusted odds ratios with corresponding 95% CIs of 10 studies on the association of gender of new-born and TIBF. The horizontal line represents the confidence interval, the box and its size in the middle of the horizontal line represents the weight of sample size. The polygon represents the pooled odds ratio. The reference category is ‘Female’. TIBF = timely initiation of breastfeeding; LIBF = late initiation of breastfeeding; REM = random-effects model.

Likewise, 13 studies^25,26,28,29,31–37,39,58^ reported the association between TIBF and ANC in 12,535 mothers (Table 1B). The pooled OR of ANC was 1.70 (95% CI 1.10 - 2.65, p = 0.02, I^2^ = 93.1%) (figure 3). Mothers who had at least one ANC visit had 70% significantly higher chance of initiating breastfeeding within one hour of birth compared with mothers who had no ANC visit. Egger’s regression test for funnel plot asymmetry was not significant (z = 0.96, p = 0.34) (Supplementary figure 2).

**Figure 3:**
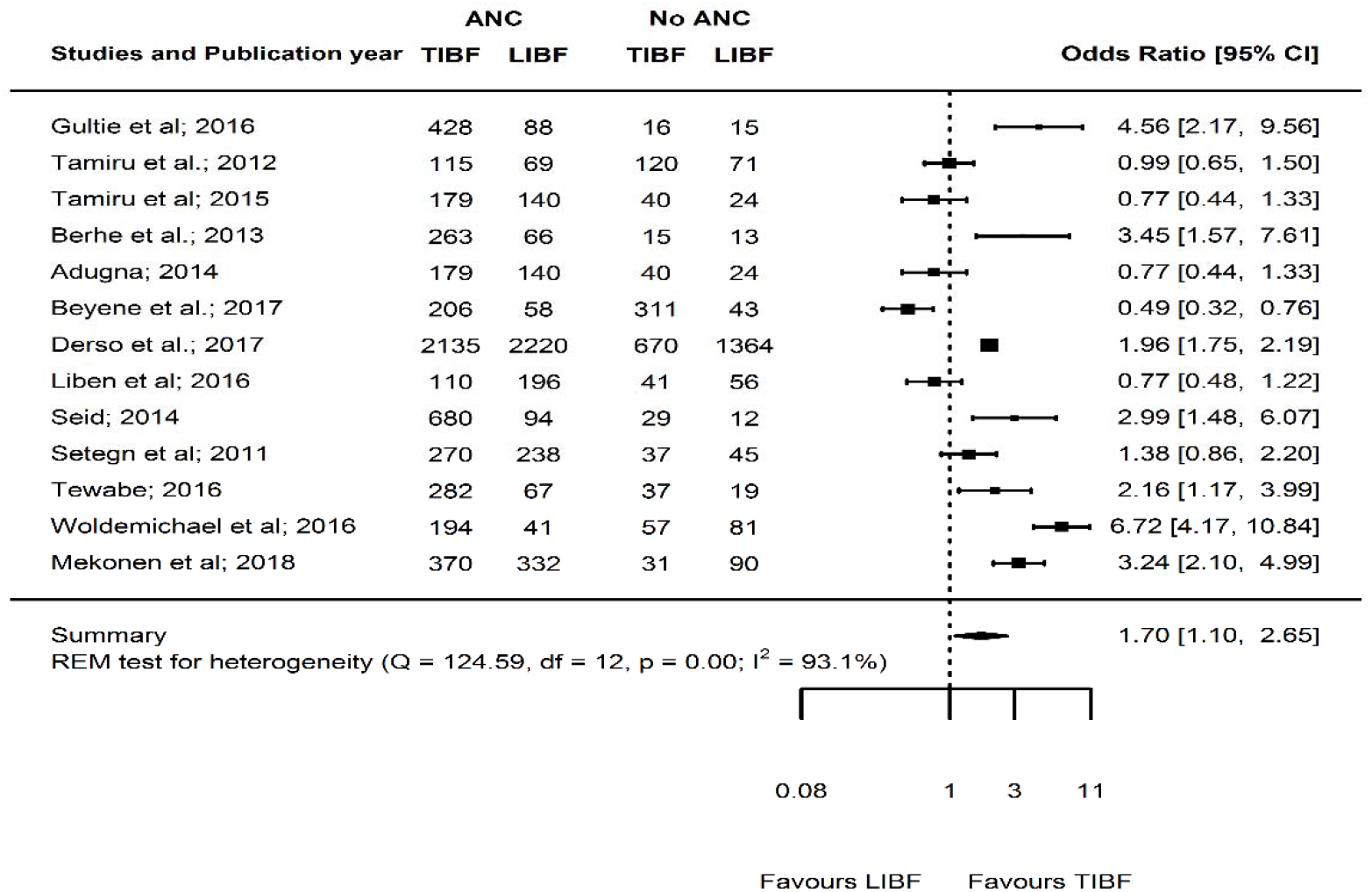
Forest plot of the unadjusted odds ratios with corresponding 95% CIs of 13 studies on the association of ANC and TIBF. The horizontal line represents the confidence interval, the box and its size in the middle of the horizontal line represents the weight of sample size. The polygon represents the pooled odds ratio. The reference category is ‘No ANC follow-up’. TIBF = timely initiation of breastfeeding; LIBF = late initiation of breastfeeding; REM = random-effects model; ANC=Antenatal care.

#### EBF

Out of the 24 studies included, 11 studies^23,24,40–48^ reported the association between EBF and gender of new-born in 6,527 mothers (Table 2A). The pooled OR of new-born gender was 1.08 (95% CI 0.86 - 1.36, p = 0.49, I^2^ = 71.7%) (figure 4). Mothers with male new-born had 8% higher chance of exclusively breastfeeding during the first six months compared with mothers with female new-born although not statistically significant. Egger’s regression test for funnel plot asymmetry was significant (z = -3.64, p < 0.001). Since significant publication bias detected, we did Duval and Tweedie trim-and-fill analysis and calculated a new effect size for gender of new-born (OR = 1.31, 95% CI 1.01 - 1.68, p = 0.04, I^2^ = 81.7%) after including imputed studies (i.e. estimated number of missing studies = 4) (Supplementary figure 3). Therefore, based on the new estimate, mothers with male new-born had 31% significantly higher chance of exclusive breastfeeding during the first six months compared with mothers with female new-born.

**Figure 4:**
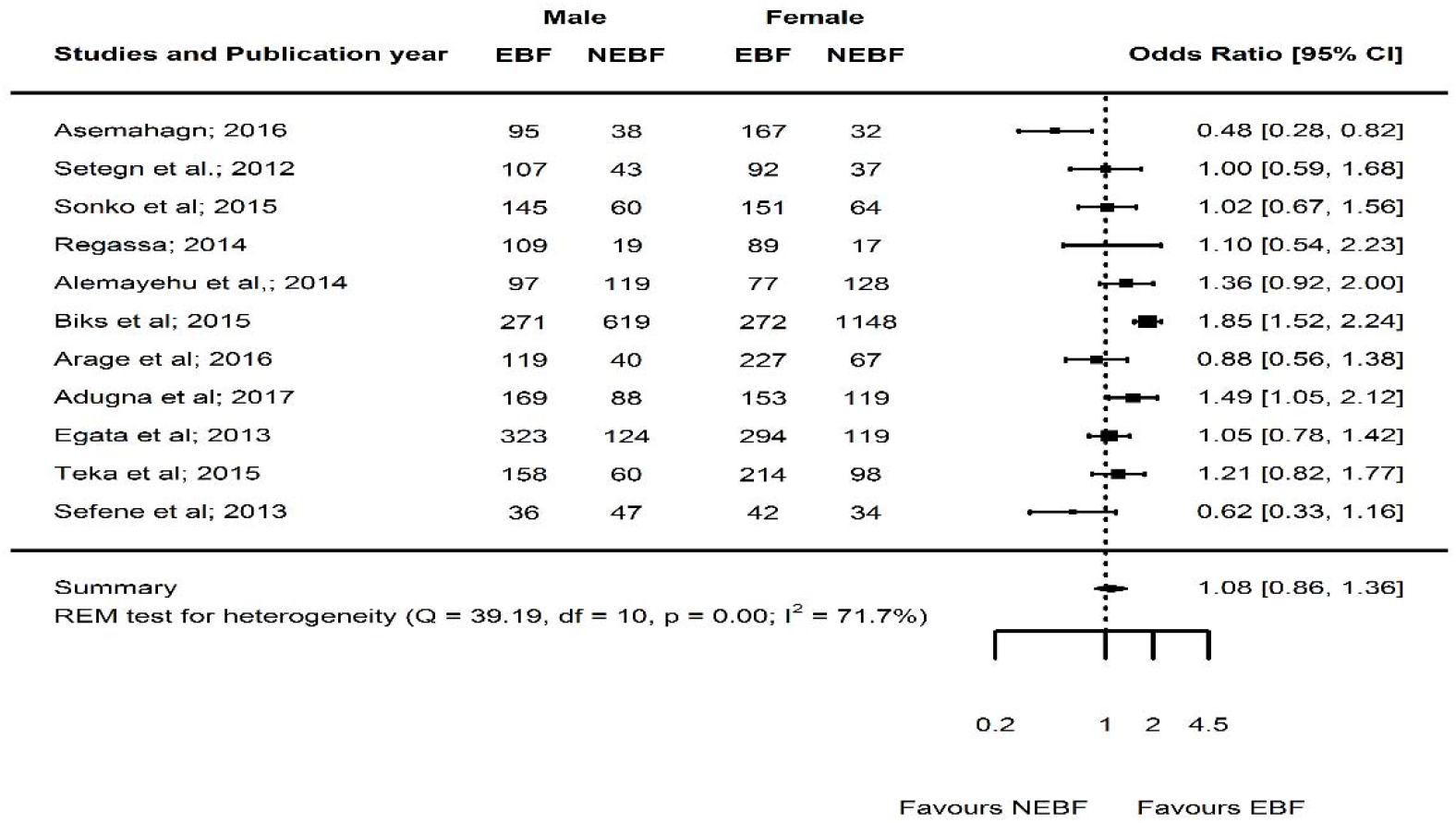
Forest plot of the unadjusted odds ratios with corresponding 95% CIs of 11 studies on the association of new-born gender and EBF. The horizontal line represents the confidence interval, the box and its size in the middle of the horizontal line represents the weight of sample size. The polygon represents the pooled odds ratio. The reference category is ‘Female’. EBF = Exclusive breastfeeding; NEBF = Non-exclusive of breastfeeding; REM = random-effects model.

Twenty-one studies^33–35,38,40–47,49–57^ reported the association between EBF and ANC in 16,052 mothers (Table 2B). The pooled OR of ANC was 2.24 (95% CI 1.65 - 3.04, p<0.0001, I^2^ = 90.9%) (figure 5). Mothers who had at least one ANC visit had 2.24 times significantly higher chance of exclusively breastfeed compared with mothers who had no ANC visit. Egger’s regression test for funnel plot asymmetry was not significant (z = 1.69, p = 0.09) (Supplementary figure 4).

**Figure 5:**
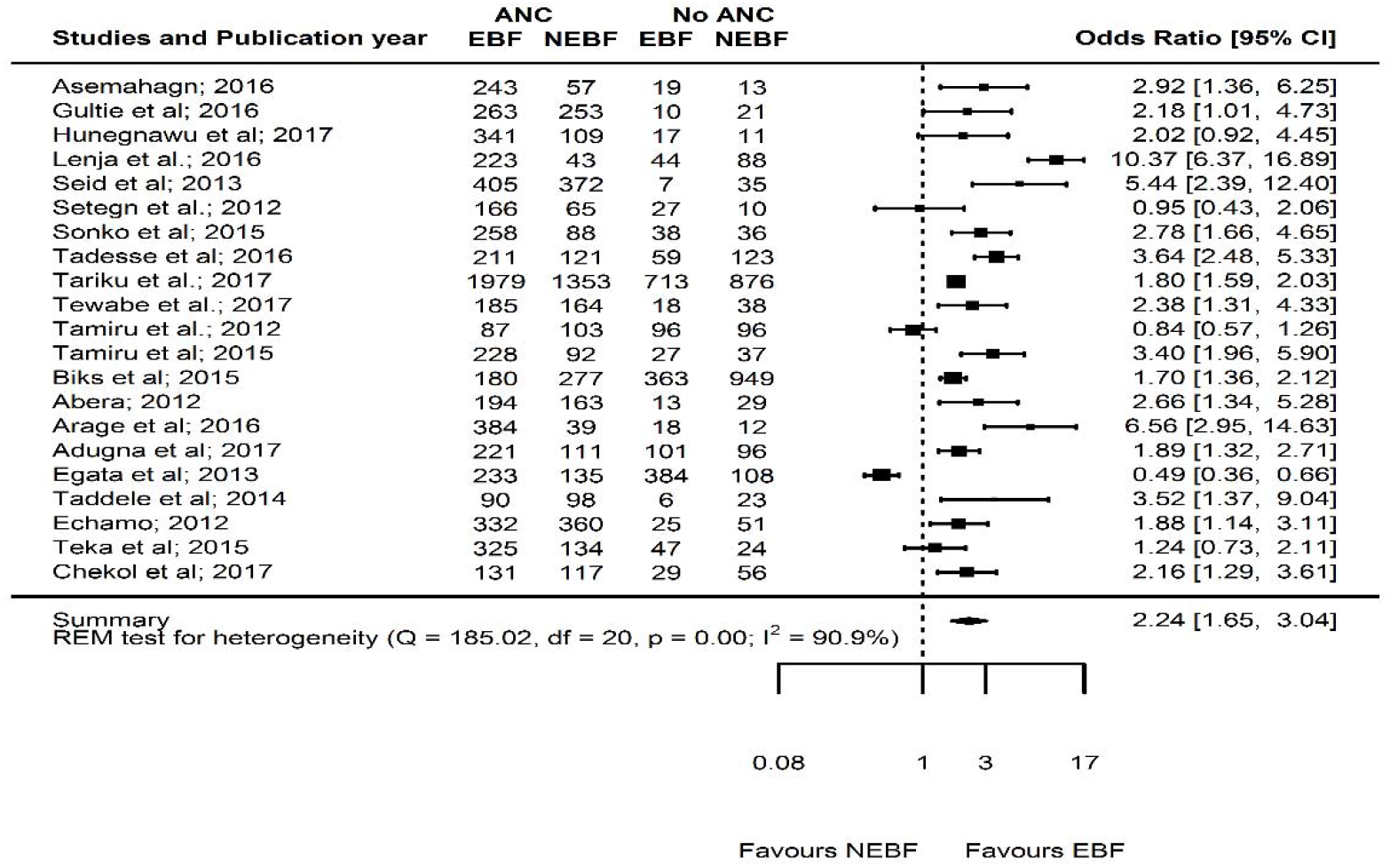
Forest plot of the unadjusted odds ratios with corresponding 95% CIs of 21 studies on the association of ANC and EBF. The horizontal line represents the confidence interval, the box and its size in the middle of the horizontal line represents the weight of sample size. The polygon represents the pooled odds ratio. The reference category is ‘No ANC follow-up’. EBF = Exclusive breastfeeding; NEBF = Non-exclusive of breastfeeding; ANC = Antenatal care; REM = random effects model.

Furthermore, seven studies^40,42,47,50,51,53,57^ reported the association between EBF and PNC in 2,995 mothers (Table 2C). The pooled OR of PNC was 1.86 (95% CI 1.41 - 2.47, p <0.0001, I^2^ = 63.4%) (figure 6). Mothers who had at least one PNC visit had 86% significantly higher chance of exclusively breastfeed during the first six months compared with mothers who had no PNC follow-up. Egger’s regression test for funnel plot asymmetry was not significant (z = -0.91, p = 0.36) (Supplementary figure 5).

**Figure 6:**
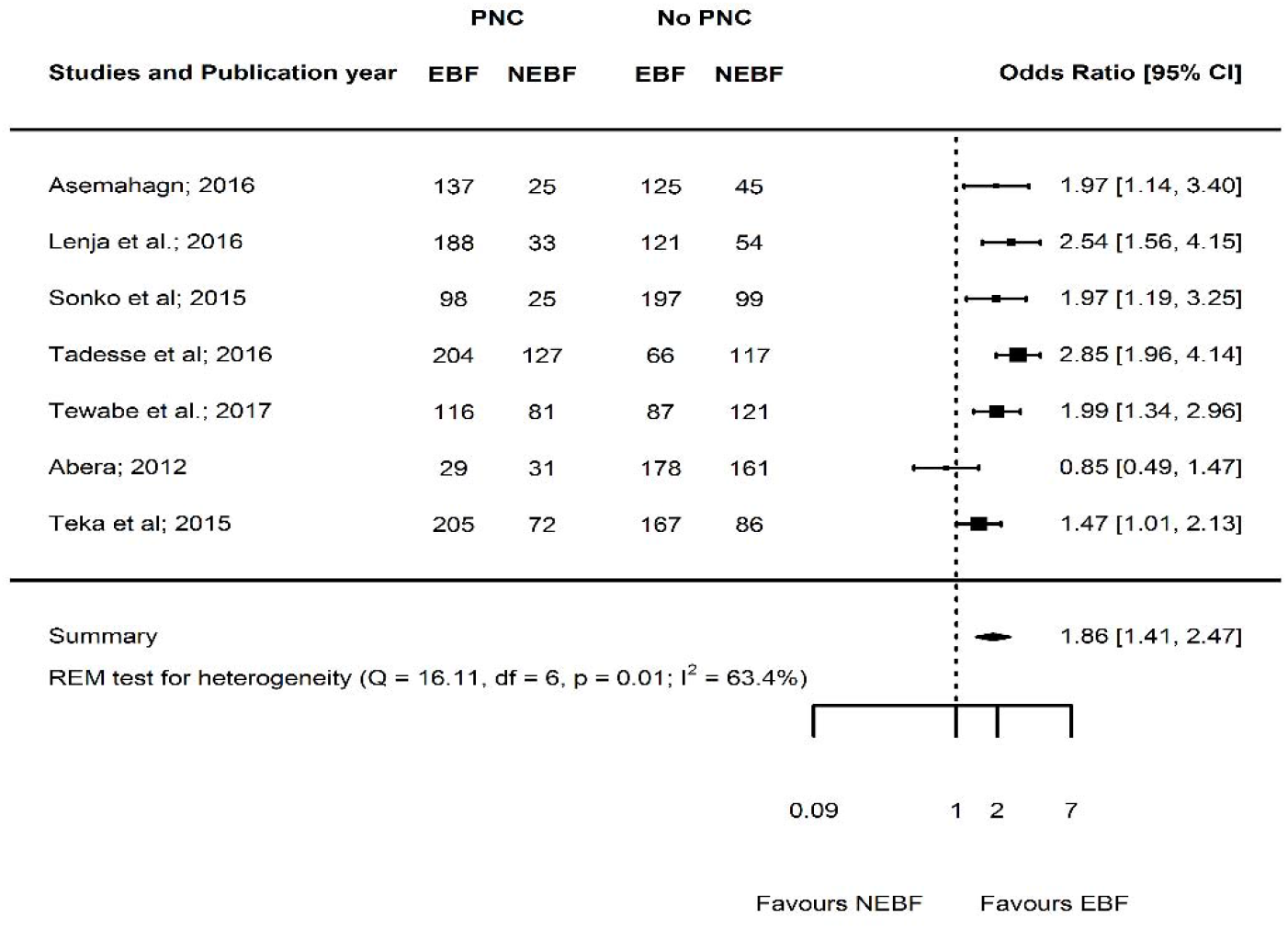
Forest plot of the unadjusted odds ratios with corresponding 95% CIs of seven studies on the association of PNC and EBF. The horizontal line represents the confidence interval, the box and its size in the middle of the horizontal line represents the weight of sample size. The polygon represents the pooled odds ratio. The reference category is ‘No PNC follow-up’. EBF = Exclusive breastfeeding; NEBF = Non-exclusive breastfeeding; PNC = Postnatal care; REM = random-effects model.

### Cumulative meta-analysis

As illustrated in figure 7, the effect of gender of new-born (figure 7) has not been changed whereas the effect of ANC on TIBF (figure 8) has been increasing over time. Similarly, the effect of gender of new-born on EBF (figure 9) has not been changed. The effect of the effect of ANC (figure 10) and PNC (figure 11) have been increasing overtime.

**Figure 7:**
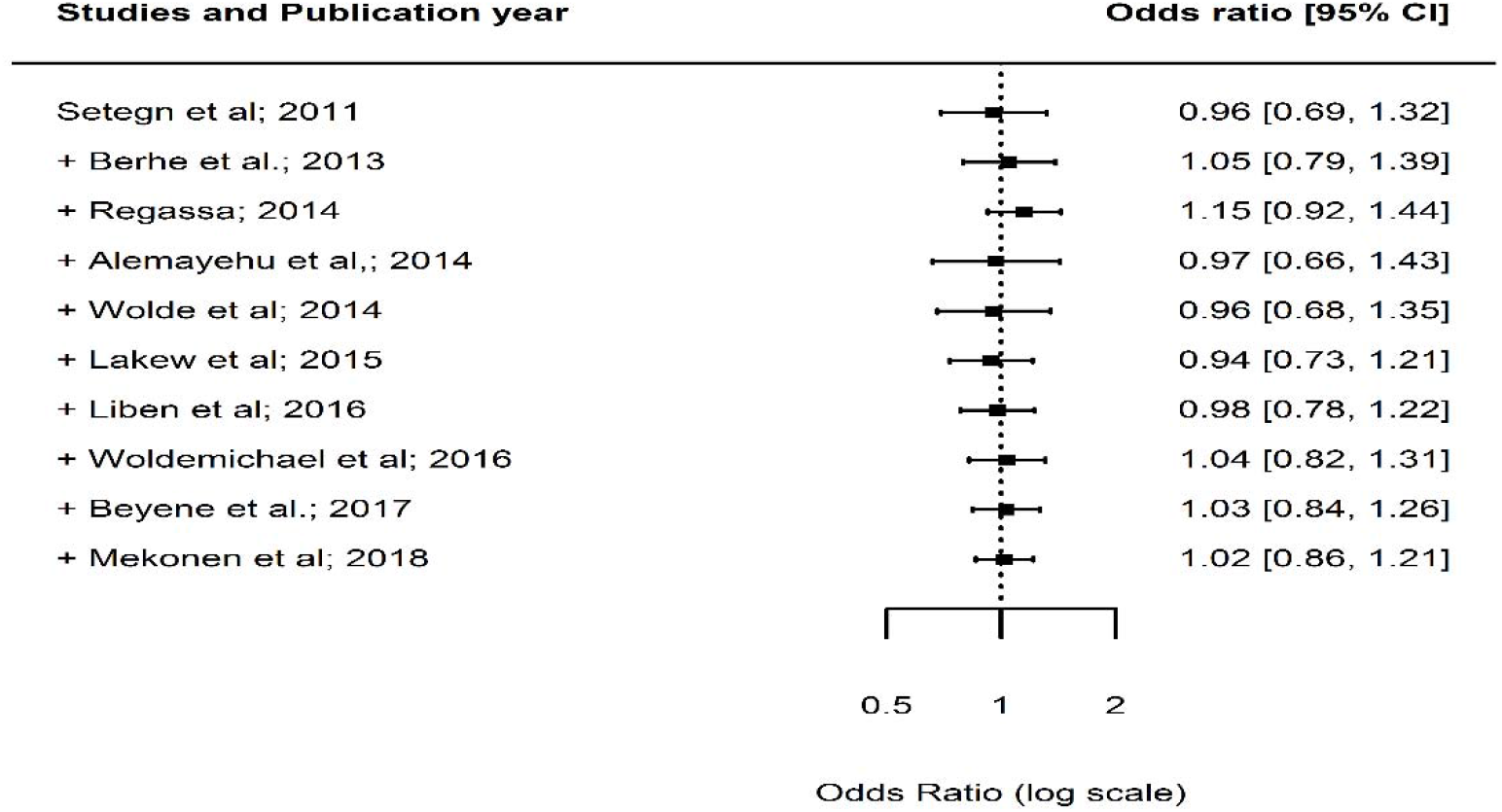
Forest plot showing the results from a cumulative meta-analysis of studies examining the effect of gender of new-born on TIBF.

**Figure 8:**
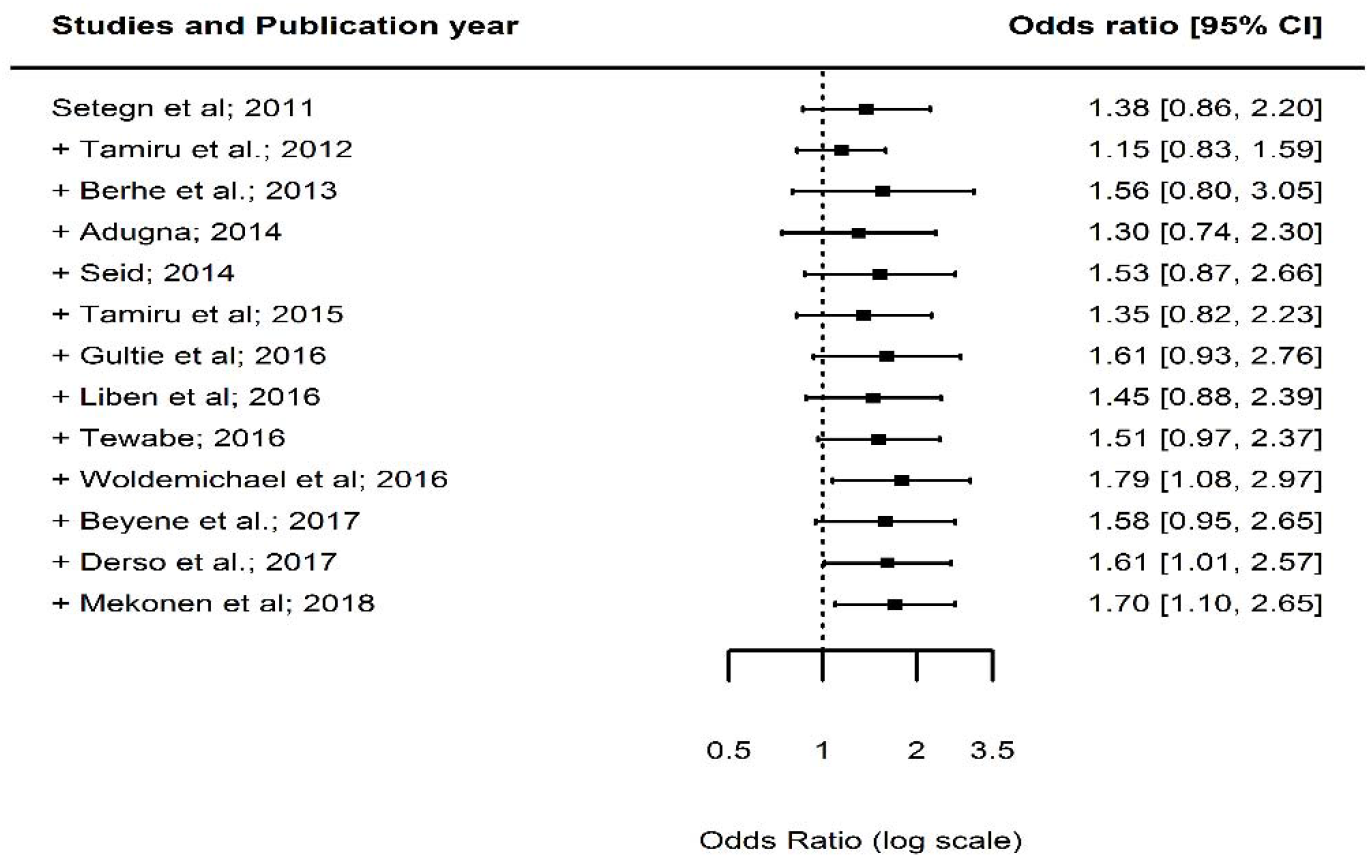
Forest plot showing the results from a cumulative meta-analysis of studies examining the effect of ANC on TIBF.

**Figure 9:**
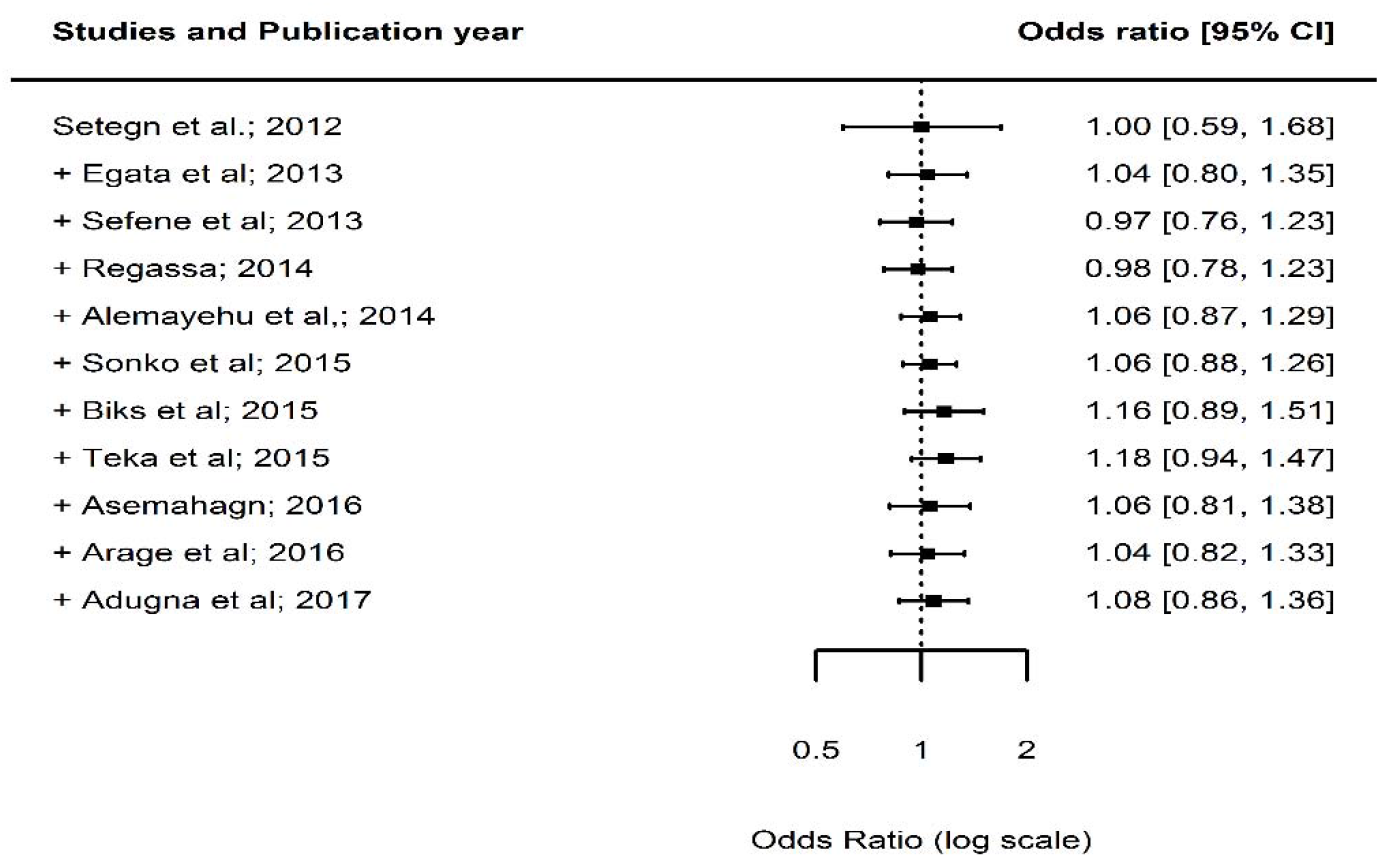
Forest plot showing the results from a cumulative meta-analysis of studies examining the effect of gender of new-born on EBF.

**Figure 10:**
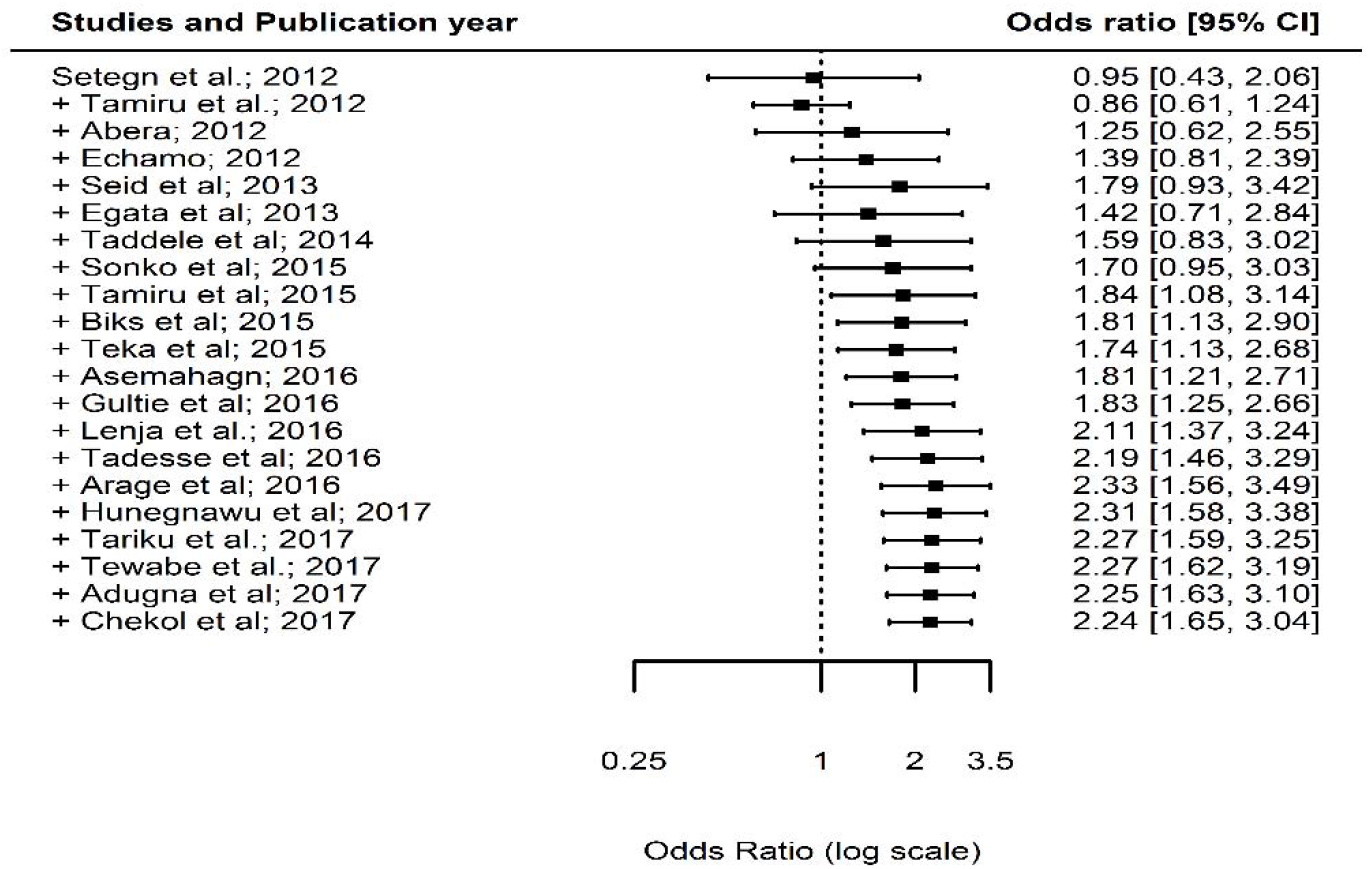
Forest plot showing the results from a cumulative meta-analysis of studies examining the effect of ANC on EBF.

**Figure 11:**
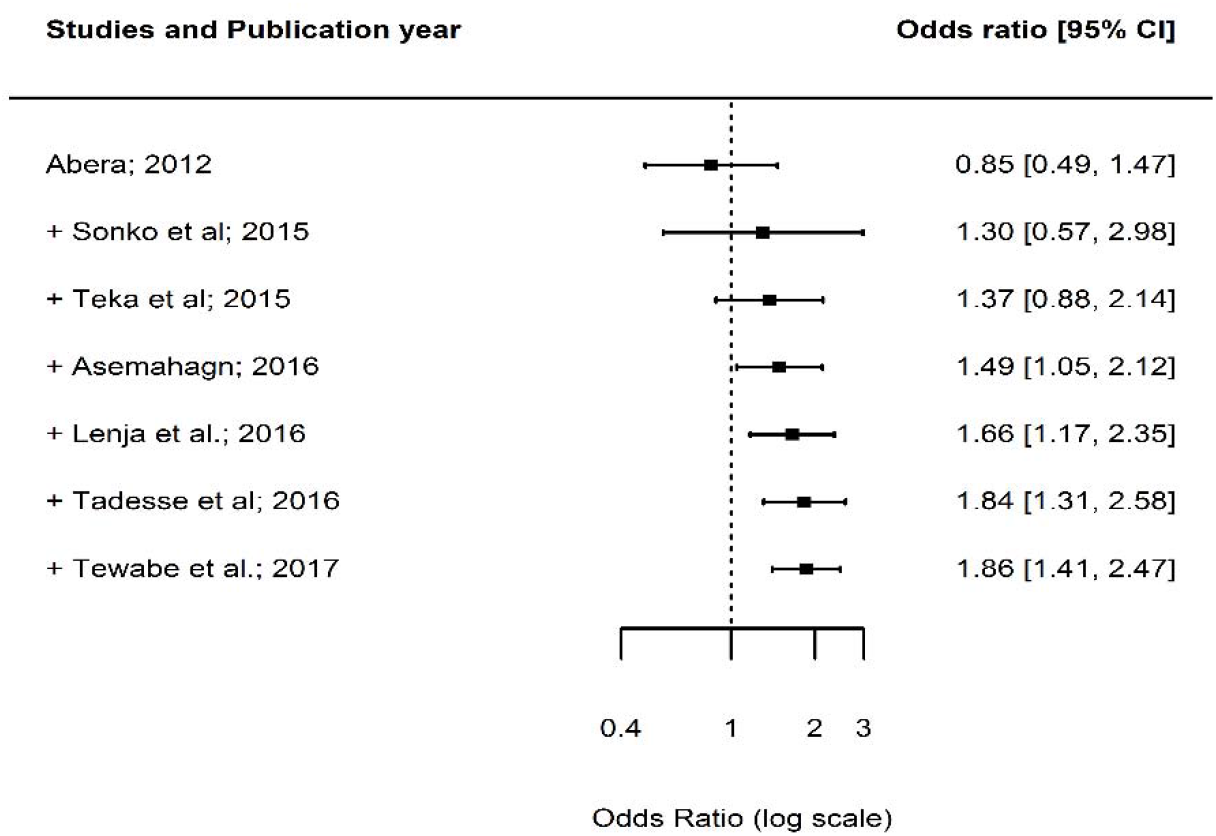
Forest plot showing the results from a cumulative meta-analysis of studies examining the effect of PNC on EBF.

### Meta-regression analysis

In studies reporting the association between TIBF and ANC, 40% of the heterogeneity was due to variation in study area (region), residence of mothers, sample size and publication year. Based on the omnibus test, however, none of these factors influenced their association (Q_M_ = 14.72, df = 8, p = 0.07). In studies reporting the association between TIBF and gender of new-born, the estimated amount of total heterogeneity was substantially low (tau^2^ = 5.4%); as a result, it is not relevant to investigate the possible reasons for heterogeneity.

In EBF, 100%, 57% and 100% of the heterogeneity among studies reporting gender of new-born, ANC and PNC were due to variation in study area (region), residence of mothers, sample size and publication year respectively. Based on the omnibus test, study area (region) and sample size positively influenced the association between gender of new-born and EBF practice (Q_M_ = 36.95, df = 7, p < 0.001). Study area (region) negatively influenced the association between ANC and EBF practice (Q_M_ = 25.75, df = 8, p = 0.001) (Table 3).

**Table 3:** Meta-regression analysis to identify possible reasons for between-study heterogeneity.

| Variables (reference category)<sup>‡</sup> | Estimate | SE | Z-value | P-value | CI.lb | CI.ub |
| --- | --- | --- | --- | --- | --- | --- |
| **TIBF** |  |  |  |  |  |  |
| **ANC** |  |  |  |  |  |  |
| Amhara region (Afar) | 1.02 | 1.01 | 1.01 | 0.31 | -0.96 | 2.99 |
| Oromia region (Afar) | 1.63 | 0.85 | 1.91 | 0.06 | -0.04 | 3.30 |
| SNNPR region (Afar) | -0.07 | 0.83 | -0.08 | 0.94 | -1.70 | 1.56 |
| Tigray region (Afar) | 1.65 | 1.23 | 1.33 | 0.18 | -0.77 | 4.07 |
| Urban residence (Rural) | 0.43 | 0.95 | 0.46 | 0.65 | -1.43 | 2.29 |
| Urban and Rural residence (Rural) | -0.08 | 0.65 | -0.12 | 0.90 | -1.35 | 1.19 |
| Sample size | -0.00 | 0.0002 | -0.31 | 0.76 | -0.0003 | 0.0002 |
| Publication year | 0.22 | 0.16 | 1.33 | 0.18 | -0.10 | 0.54 |
| **EBF** |  |  |  |  |  |  |
| ***Gender of new-born*** |  |  |  |  |  |  |
| Oromia region (Amhara) | 0.05 | 0.26 | 0.19 | 0.85 | -0.45 | 0.55 |
| SNNPR region (Amhara) | 0.72 | 0.24 | 2.96 | 0.003 | 0.24 | 1.19 |
| Tigray region (Amhara) | 0.55 | 0.20 | 2.71 | 0.01 | 0.15 | 0.94 |
| Urban residence (Rural) | 0.40 | 0.31 | 1.28 | 0.19 | -0.21 | 1.02 |
| Urban and Rural residence (Rural) | 0.24 | 0.31 | 0.76 | 0.45 | -0.37 | 0.85 |
| Sample size | 0.001 | 0.0002 | 4.77 | <0.0001 | 0.0005 | 0.001 |
| Publication year | -0.07 | 0.09 | -0.68 | 0.49 | -0.26 | 0.13 |
| ***ANC*** |  |  |  |  |  |  |
| Harari region (Amhara) | -0.21 | 0.69 | -0.31 | 0.76 | -1.58 | 1.15 |
| Oromia region (Amhara) | -1.62 | 0.51 | -3.18 | 0.002 | -2.62 | -0.62 |
| SNNPR region (Amhara) | -0.12 | 0.33 | -0.36 | 0.72 | -0.77 | 0.53 |
| Tigray region (Amhara) | -0.77 | 0.63 | -1.21 | 0.23 | -2.01 | 0.47 |
| Urban residence (Rural) | -0.41 | 0.33 | -1.25 | 0.21 | -1.04 | 0.23 |
| Urban and Rural residence (Rural) | -0.47 | 0.44 | -1.05 | 0.29 | -1.34 | 0.40 |
| Sample size | -0.0001 | 0.0001 | -0.66 | 0.51 | -0.0004 | 0.0002 |
| Publication year | -0.04 | 0.09 | -0.47 | 0.64 | -0.23 | 0.14 |
| ***PNC\**** |  |  |  |  |  |  |
| Harari region (Amhara) | -0.25 | 1.30 | -0.19 | 0.85 | -2.79 | 2.29 |
| SNNPR region (Amhara) | 0.28 | 0.44 | 0.65 | 0.52 | -0.56 | 1.13 |
| Tigray region (Amhara) | -0.19 | 0.67 | -0.28 | 0.78 | -1.49 | 1.12 |
| Sample size | 0.001 | 0.002 | 0.41 | 0.68 | -0.003 | 0.004 |
| Publication year | 0.13 | 0.27 | 0.49 | 0.62 | -0.39 | 0.66 |

<sup>¥</sup> = Since we do not have a specific hypothesis, the reference category is selected arbitrarily; \* = Residence is dropped from the model due to small sample size of included studies; CI.lb = Confidence interval, lower bound; CI.ub = Confidence interval, upper bound; SNNPR = Southern Nations, Nationalities and Peoples' Region.

## Discussion

This meta-analysis assessed the association of timely initiation of breastfeeding (TIBF) and exclusive breastfeeding (EBF) with gender of new-born, antenatal (ANC) and postnatal care (PNC). The key findings were (1) ANC, PNC and gender of new-born significantly associated with EBF and (2) ANC significantly associated with TIBF but not gender of new-born.

In congruent with our hypothesis and the large body of global evidence,^59–64^ our finding indicated that mothers who had at least one antenatal visit had significantly higher chance of initiating breastfeeding within one hour of birth and exclusively breastfeed for the first six months compared with mothers who had no ANC visit. This may be due to the fact that health professionals provide breastfeeding guidance and counseling/health education during ANC visit. This justification is supported by our previous meta-analysis^7^ and WHO/UNICEF presumption which emphasizes promoting breastfeeding during pregnancy through the Baby-Friendly Hospital Initiative (BFHI) program. Ethiopia has also adopted BFHI as part of the national nutrition program and is now actively working to integrate to all public and private health facilities and improving breastfeeding practice as a result.

We also showed that mothers who had at least one PNC visit had nearly twice higher chance of exclusively breastfeeding during the first six months compared with mothers who had no PNC follow-up; this result supported our hypothesis. Similarly, several studies have reported a significantly high rate of EBF in mothers who had a postnatal visit at health institution^64^ or postnatal home visit.^65^ The possible justification could be that postnatal visit health education may positively influence the belief and decision of the mothers to exclusively breastfeed. Previous studies have also shown that postnatal education and counseling are important to increase EBF.^66^ In addition, in our previous meta-analyses, we showed that guidance and counseling during ANC or PNC significantly associated with high rate EBF.^7^ Furthermore, postnatal care may ease breastfeeding difficulty, increase maternal confidence and encourage social/family support which lead the mother to continue EBF for 6 months.

Finally, in agreement with previous studies^67–69^ and our hypothesis, we uncovered gender of new-born significantly associated with EBF. Mothers with male new-born had 31% significantly higher chance of exclusively breastfeeding during the first six months compared with mothers with female new-born. This meta-analysis result disproved the traditional perception and believe in Ethiopia that male new-born have pre-lacteal feeding to be strong and healthy compared with female new-born; however, further investigation is required. On the other hand, we showed that gender of new-born not significantly associated with TIBF. Several studies^61,64^ also showed that gender of new-born is not significantly associated with breastfeeding practice. This discrepancy across studies may be due to the socio-cultural difference. In addition, the non-significant association between gender of new-born and TIBF may be due to lack of adequate power given that only 10 studies were used for pooling the effect size.

This systematic review and meta-analysis was conducted based on the registered and published protocol,^13^ and Preferred Reporting Items for Systematic Reviews and Meta-Analyses (PRISMA) guidelines for literature reviews. In addition, publication bias was quantified using Egger’s regression statistical test and NOS was used to assess the quality of studies. Since it is the first study in Ethiopia, the information could be helpful for future researchers, public health practitioners, and healthcare policymakers. The inclusion of large sample size and recent studies are further strengths of this study. This study has limitations as well. Almost all included studies were observational which hinder causality inference. Even though we have used broad search strategies, the possibility of missing relevant studies cannot be fully exempted and the finding may not be nationally representative. Based on the conventional methods of the heterogeneity test, a few analyses suffer from high between-study variation. The course of heterogeneity was carefully explored using meta-regression analysis and this variation may be due to the difference of study area (region), residence of mothers, sample size, publication year or other residual factors; therefore, the result should be interpreted with caution. Moreover, the dose-response relationship between the number of ANC/PNC visits and breastfeeding practices was not examined. Lastly, a significant publication bias was detected in studies reported the association between EBF and gender of new-born. We did Duval and Tweedie trim-and-fill analysis to adjust publication bias and to provide an unbiased estimate; however, the result should be interpreted cautiously.

## Conclusions

We found, in line with our hypothesis, gender of new-born, ANC and PNC significantly associated with EBF. Likewise, ANC significantly associated with TIBF. This meta-analysis study provided evidence on breastfeeding practices and its associated factors in an Ethiopian context, which can be useful for cross-country and cross-cultural comparison and for breastfeeding improvement initiative in Ethiopia. Most importantly, this study provides an overview of up-to-date evidence for public nutrition professionals and policymakers. In addition, the result indicates that increasing the utilization of antenatal and postnatal care have a positive effect on breastfeeding practices. This signifies stakeholders would provide emphasis on ANC and PNC service to achieve WHO breastfeeding goal. From the research point of view, in general, intervention- and outcome-based studies of breastfeeding in Ethiopia are required.

## Data sharing statement

All data generated or analysed in this study are included in the article and its supplementary files.

## Competing interests

The authors declare that they have no competing interests.

## Funding

Not applicable

## Authors Contribution

NT and TD conceived and designed the study. TD developed a syntax for searching databases, analyzed the data and interpreted the results. TD and SM wrote and revised the manuscript. All authors read and approved the final manuscript.

## Acknowledgment

Our special gratitude forwarded to Sjoukje van der Werf (University of Groningen, the Netherlands) for her support to develop the search strings and Balewgizie Sileshi (University of Groningen, the Netherlands) for his support during the title and abstract screening.

